# A simplified spiking model of grid-cell scale and intrinsic frequencies

**DOI:** 10.1101/544882

**Authors:** Diogo Santos-Pata, Riccardo Zucca, Héctor López-Carral, Paul F. M. J. Verschure

**Author notes:** Corresponding author, Email address (Paul F. M. J. Verschure).

## Abstract

The hexagonal tessellation pattern of grid cells scales up progressively along the dorsal-to-ventral axis of the medial entorhinal cortex (MEC) layer II. This scaling gradient has been hypothesized to originate either from inter population synaptic dynamics as postulated by attractor networks, from projected theta frequencies to different axis levels, as in oscillatory models, or from cellular dynamics dependent on hyperpolarization-activated cation currents. To test the hypothesis that intrinsic cellular properties account for the scale gradient as well as the different oscillatory frequencies observed along the dorsal-to-ventral axis, we have modeled and analyzed data from a population of grid cells simulated with spiking neurons interacting through low-dimensional attractor dynamics. To investigate the causal relationship between oscillatory frequencies and grid scale increase, we analyzed the dominant frequencies of the membrane potential for cells with distinct after-spike dynamics. We observed that intrinsic neuronal membrane properties of simulated cells could induce an increase of grid scale when modulated by after-spike reset values. Differences in the membrane potential oscillatory frequency were observed along the simulated dorsal-to-ventral axis, suggesting that, rather than driving to the increase of grid scale as proposed by interference models of grid cells, they are the result of intrinsic cellular properties of neurons at each axis level. Overall, our results suggest that the after-spike dynamics of cation currents may play a major role in determining the grid cells’ scale and that oscillatory frequencies are a consequence of intrinsic cellular properties that are specific to different levels of the dorsal-to-ventral axis in the MEC layer II.

## 1. Introduction

Grid cells found in layer II of the medial entorhinal cortex (MEC) present multiple regularly-spaced firing fields organized in a triangular tessellation that spans the entire explored environment [1, 2]. Functionally, grid cells represent a spatial metric system signaling the position of the animal in the environment. Together with sensory cells in the lateral entorhinal cortex (LEC), grid cells in MEC layer II project to both the dentate gyrus (DG) and CA3 neurons of the hippocampus proper [3, 4]. Thus, the mammalian hippocampus robustly encodes spatial representation using a combination of environment-related spatial and sensory information.

Since the discovery of grid cells, several computational models have been proposed to describe the spatial and temporal properties of grid fields’ formation. Most of the proposed models can be categorized into two groups: oscillatory interference [5] and attractor dynamics [6, 7]. For the former group, the hexagonal grid pattern emerges from the interaction of multiple phase-synchronized oscillations that are based on the animal’s speed vector projected to MEC layer II from earlier MEC layers. Thus, at the computational level, manipulating the amplitudes and phase differences of these oscillations would modulate the scale of the resulting grid cells. On the other hand, in the attractor-based models, the distribution of synaptic weights within an all-to-all network creates a characteristic “bump” of activity that converges to stable attractor points.

The network weights configuration is updated according to the spatial motion of the agent at every time (*t*), which allows for the characteristic periodic firing across the explored environment. Recent studies on the intrinsic cellular properties of grid cells support the idea of low-dimensional continuous attractor dynamics in the grid cells’ system, favoring the computational principles of attractor-based models of grid cells [8, 9].

The size and spacing of grid cells’ firing fields have been shown to increase progressively along the dorsal-to-ventral axis of the MEC [1, 10, 11]. Functionally, such a scale gradient has been suggested to operate as an accurate path-integration mechanism projecting to the DG and CA3 hippocampal sub-regions [10]. Moreover, the interaction between grid scales and other spatially tuned cells have been suggested to serve for minimizing errors in path integration [12].

Despite observations on hyperpolarization-activated cyclic nucleotide-gated (HCN) channels’ disruption and its effects on grid scale [13], the mechanism underlying the differences in scale of such neural populations is still not clear. Different sub-threshold theta oscillatory frequencies have been measured in vitro in neurons along the dorsal-to-ventral axis, suggesting that individual cells’ intrinsic frequencies might play a key role on grid cells’ scale [14]. Moreover, it has been shown that the distance from the dorsal surface is accompanied by a decrease in oscillatory frequency in MEC layer II [15].

From the continuous attractor model perspective, different scales are often obtained by manipulating the variance of the Gaussian synaptic distributions. However, given recent insight on the effects of HCN channels disruption in grid cells’ metrics, the distribution of synaptic weights might not be the main factor accounting for grid scale and stability of the network activity.

Coherent with such idea, previous computational models of grid cells [16] have explicitly pointed out that differences in grid cells’ scale along the dorsal-to-ventral axis are linked to differences in the cells’ intrinsic frequencies.

Indeed, a systematic topographical change in time constants of hyperpolarization-activated cation currents (*I*_*h*_) of stellate cells has been observed in vitro [14]. Moreover, those topographical changes correlate with membrane potential oscillation frequency and differences in the time constant of the sag response.

This suggests that different *I*_*h*_ kinetics, which are regulated by the HCN family proteins, may play a critical role in the change of oscillatory frequencies along the dorsal-to-ventral axis and the topographical expansion of grid scale [15]. Forebrain-specific knockout of the HCN1 subcomponent in mice has been shown to selectively affect the Y-intercept of the grid scale, indicating that those elements of the HCN family are involved in grid scale modulation [13, 17].

Previous studies have addressed the question of how intrinsic cell’s frequency affects the grid scale along the dorsal-to-ventral axis [18]. Specifically, they proposed a model where the addition of physiologically plausible after-spike dynamics modulates the observed increase in grid scale along the dorsal-ventral axis of MEC.

Whether the membrane potential oscillatory frequency is sufficient to determine the grid scale is still unclear. In attractor models of grid cells’ formation, the scale of the grid is modulated by a gain parameter affecting the synaptic connectivity of the network and thus the speed at which the activity bump moves along the network as well [18, 19]. However, there is no biological evidence for such connectivity matrix discretization. In interference models, the differences in grid scale are generated due to amplitude and phase changes in the oscillatory inputs to the grid cells network. Despite the fact that differences in the oscillatory frequencies are observable in biological systems, it is not clear whether it emerges from intrinsic or extrinsic network dynamics. The fact that knocking out HCN family type genes disrupts the normal progressive scale increase raises the question as to whether such scale gradient is a network or a cellular property. We address this by presenting a simplified spiking computational model that describes the generation of the spatial and temporal properties of grid cells found in physiological studies.

## 2. Materials and methods

In order to explore the effects of intrinsic cellular properties on differences in spatial grid scale found in the dorsal-to-ventral axis of MEC layer II and the impact on the oscillatory frequency at each axis level, we created a simulated environment where a virtual agent was randomly exploring either a 1D linear track or a 2D square arena (see Fig. 1). In both environments, the agent’s speed vector was fed into an ensemble of simulated neuronal populations (described below). Keeping in line with the findings from Yoon et al. [9] regarding the evidence of low-continuous attractor dynamics in grid cells’ populations, we built on elements of a previously presented grid cells model based on attractor dynamics [19] and translated it to a spiking neuronal model, approximating the spiking behavior of MEC layer II stellate cells. At the topological level, the network is based on the twisted toroidal architecture and synaptic weights are dependent on the Cartesian distance of each cell to its postsynaptic cells and updated according to the speed vector of the simulated agent moving within the virtual environment (see [19] for more details).

**Figure 1:**
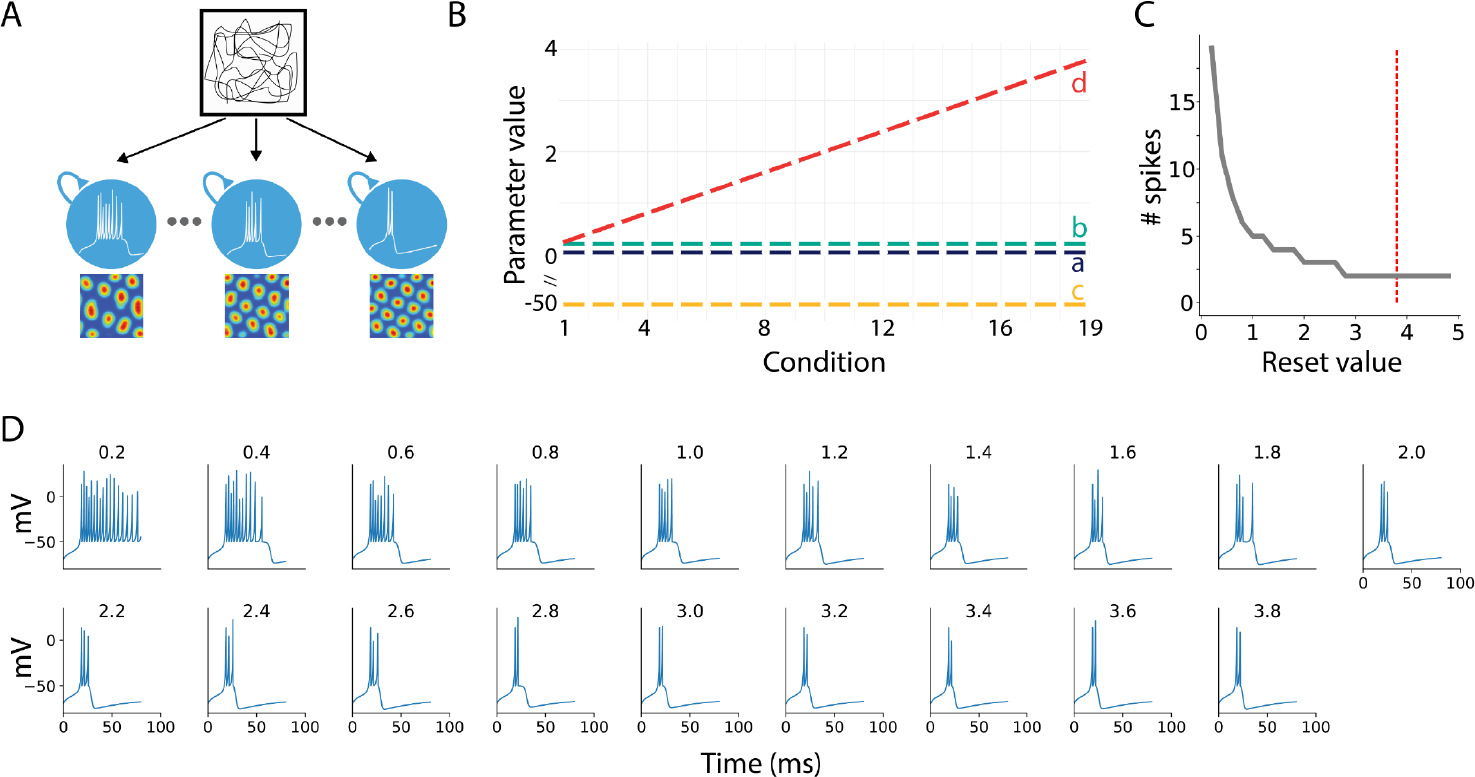
Illustrative description of the methods used in this study. **A**. A virtual agent is set to randomly explore a squared virtual arena. During exploration, 19 populations of grid cells are activated and all receive inputs from the vestibular system encoding the speed vector of the virtual agent. Each population of grid cells was initialized with specific cell model parameters as shown in B. The spatial rate map of each cell belonging to each population was stored for further analysis. **B**. Izhikevich model parameters used per each population (condition). Every parameter value was kept constant for every population, with the exception of the *d* parameter ranging from 0.2 to 3.8 in steps of 0.2. **C**. Number of spikes as a function of the *d* parameter. With all the other cell’s parameters maintained constant a plateau is observed for reset values larger than 3. **D**. Simulations of hippocampal stellate neurons for the 19 populations included in the model by varying the *d* parameter.

A total of 19 populations of grid cells were created, each containing 100 neurons connected in an all-to-all fashion (Fig. 1A). The model was implemented in the NEST neural networks simulator [20], and all the analyses were done using the SciPy python scientific library [21]. The simulations used the simplified Izhikevich’s spiking model [22], which allows for the direct manipulation of the resonant properties affected by HCN channels in biological stellate cells [13] through a single parameter. In the model, parameters were tuned to reproduce the bursting behavior of MEC layer II grid cells observed experimentally in stellate cells recordings [22, 23, 24]. The activation function of each neuron in the network was defined by a system of ordinary differential equations given by:

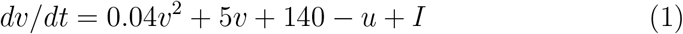

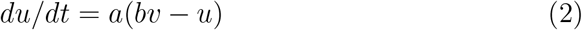

where 0.04*v*^2^ + 5*v* + 140 mimics the spike initiation dynamics of a neuron, *I* represents synaptic currents or injected DC-currents, *v* represents the cell’s membrane potential and *u* describes the membrane recovery variable. *a* describes the time scale of the recovery variable *u* and *b* describes the sensitivity of the recovery variable *u* to the subthreshold fluctuations of the membrane potential *v*.

The after-spike resetting mechanism is given by:

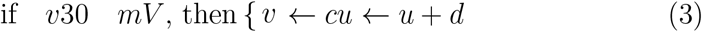

where *c* and *d* describe the after-spike membrane value and recovery variable, respectively.

To test whether the modulation of HCN channels is sufficient to trigger changes in grid cells’ scale, the after-spike reset value of cation currents *d* was varied across populations in the range 0.2 - 3.8 mV/ms with linearly increasing steps (Fig. 1B). The parametric space was defined in order to maintain the spiking behavior of stellate cells (Fig. 1C-D). All the other model parameters were kept constant over all the neuronal populations.

The parameter *a*, which describes the cell’s current recovery variable time scale, was set to 0.03. The parameter *b*, describing the cell’s current recovery, was set to 0.2. The parameter *c*, describing the after-spike reset value of the cell’s membrane potential, was set to −50 mV. The spike train activity of each cell was recorded and used for the subsequent analysis.

The virtual agent’s method of exploration was set to exhibit two different behaviors depending on the environment. In order to analyze differences in periodicity and size of grid cells’ firing fields for populations with different *I*_*h*_ currents, the first behavior of the agent was to run back-and-forth in a linear track environment. For the second environment, the square arena, the agent would explore the arena randomly. Thus, in the second environment, the characteristic 2D rate maps of grid cells can be depicted.

Grid cell’s activity was initialized with random activity between 0 and 1*/N* (number of neurons in each grid-cell module) and was modulated by the speed vector of the agent’s translation at each time step.

The network’s input is reliant on the speed vector of the simulated agent’s exploration, *s* ≔ (*s*_*x*_, *s*_*y*_), allowing the activity bump of the network shifts along the neural sheet when the agent moves according to its speed vector. Whereas in the original grid cell model proposed by [19] the speed vector *s* is susceptible of modulation via a gain parameter, affecting the grid scale, in our simulations, the speed vector was kept constant throughout the simulated conditions.

The synaptic weights distribution was defined as:

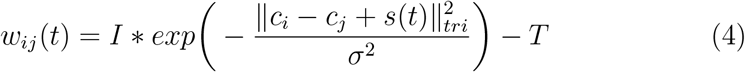

where 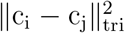 denotes the Cartesian distance between cells *c*_*i*_ and *c*_*j*_ in the network matrix, I (= 0.3) defines the synaptic strength, *σ* (= 0.48) modulates the width of the synaptic weight distribution and T (= 0.05) is the excitatory and inhibitory distance threshold.

### 2.1. Data analysis

Occupancy maps were calculated as the total time an agent spent in each spatial bin (50 × 50 pixels) within the virtual arena. Rate maps were then obtained by normalizing each cell’s spiking activity within a spatial bin with the agent’s occupancy map. Autocorrelograms were then obtained by the spatial autocorrelation of the rate map of each cell in the 2-dimensional plane.

Frequency analyses were obtained by averaging the dominant frequency provided by the power spectral density (PSD). To compute the dominant membrane potential oscillation frequency, continuous, contiguous and non-overlapping windows of 10 seconds were extracted from each cell membrane potential and their PSD was computed. The second highest peak of the averaged PSD was considered the dominant frequency for a given cell.

## 3. Results

To verify that the manipulation of the intrinsic cellular properties in the chosen cell model simulation would not affect the attractor mechanism of the networks, we set every simulation to be a random state of activity and visually ensured that an activation bump was formed and remained stable throughout the virtual agent navigation. The formation of the activity bump during the initial simulation steps for three representative populations with different after-spike reset values are shown in Fig. S1.

### 3.1. Grid scale is modulated by I_h_

#### 3.1.1. Linear track simulation

The virtual agent was set to run along a linear track environment, measuring 200 virtual units, at a constant speed of 20 virtual units/second. A total of 1900 spike trains were recorded.

To test whether the modulation of HCN channels is sufficient to trigger changes in grid cells’ scale, the after-spike reset values of cation currents *d* were varied across populations. Fig. 2 illustrates the effects of varying the after-spike dynamics on the firing fields of 4 representative cells from the simulated dorsal-to-ventral axis level conditions (*d* = 3.6, 2.2, 1.2, 0.2). Within the linear track, spike activity relative to the agent’s position in the environment points to an increase in firing-fields size and distance that is dependent on the value of *d*. The effect appears more evident from the spike density plots (Fig. 2 middle subplots) and by the periodic regions of high activity observed in the spike density autocorrelations (Fig. 2 bottom subplots).

**Figure 2:**
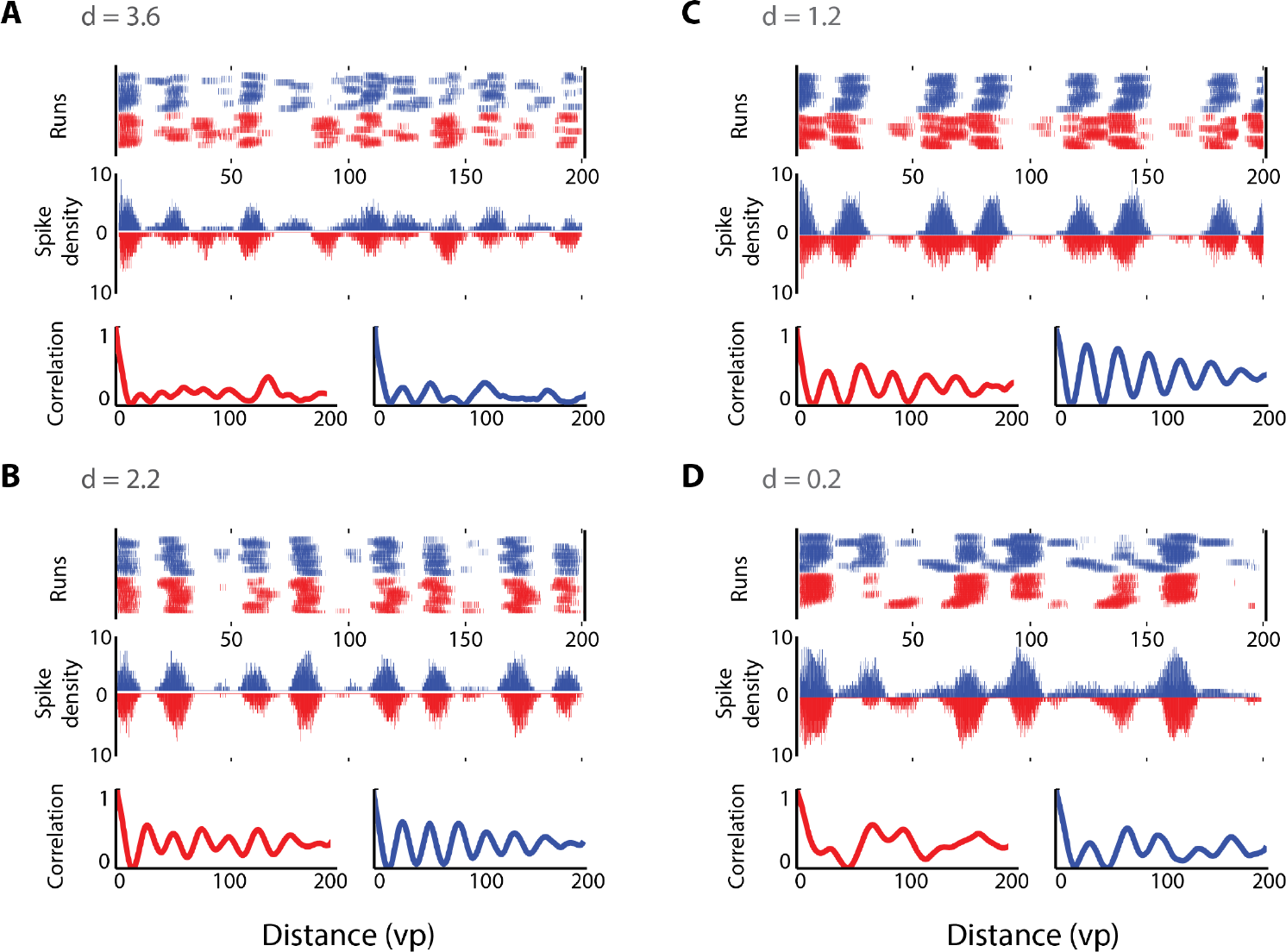
Progressive increase in grid scale from dorsal to ventral MEC. Spike-trains of four representative cells from dorsal (A) to ventral (D) axis populations. Left trajectories (red) and right trajectories (blue) are differentiated. Top subplots represent the raw spikes against the position of the linear track per each run. Middle subplots represent the spike density per position (same color code as in top plot). Bottom subplots show the correlations of spike density along the linear track. Higher to lower oscillation of spike density and correlation is observed from dorsal to ventral levels.

To quantify the increase in grid scale at the population level, we have quantified each simulated cell firing field’s size and distance (Fig. 3). To do so, we obtained the firing rate, spike count, at each position of the linear track as in Fig. 2 (bin size = 5 virtual points). A peak detection algorithm was applied to identify the firing fields. Firing fields whose peak rate was larger than 1.2 standard deviations (z-scored) were included in the sample and the averaged distance between consecutive fields of each cell was computed. Increasing the hyperpolarization reset value *d* caused a decrease in the averaged firing field distance (Pearson *r* = −0.34, *p* < 0.01, Fig. 3-top left). Similarly, in order to quantify each cell’s firing field size, we obtained the spatial distance between the two firing rate points on each side of its peak whose first derivative was ≥ 0. As for distance, we found that firing fields size is negatively modulated by the after-spike reset parameter (Pearson *r* = −0.31, *p* < 0.01, Fig. 3-top right). We next asked whether firing fields size and distance were equally affected by the hyperpolarization reset value of each condition. To do so, we averaged firing fields size and distance of cells belonging to the same condition (Fig. 3-bottom). A strong correlation between size and distance revealed significance (Pearson *r* = 0.79, *p* < 0.01). Moreover, the increase in these spatial measures was accompanied by a decrease in the hyperpolarization reset value (see *I*_*h*_ condition in Fig. 3-bottom, colorbar).

**Figure 3:**
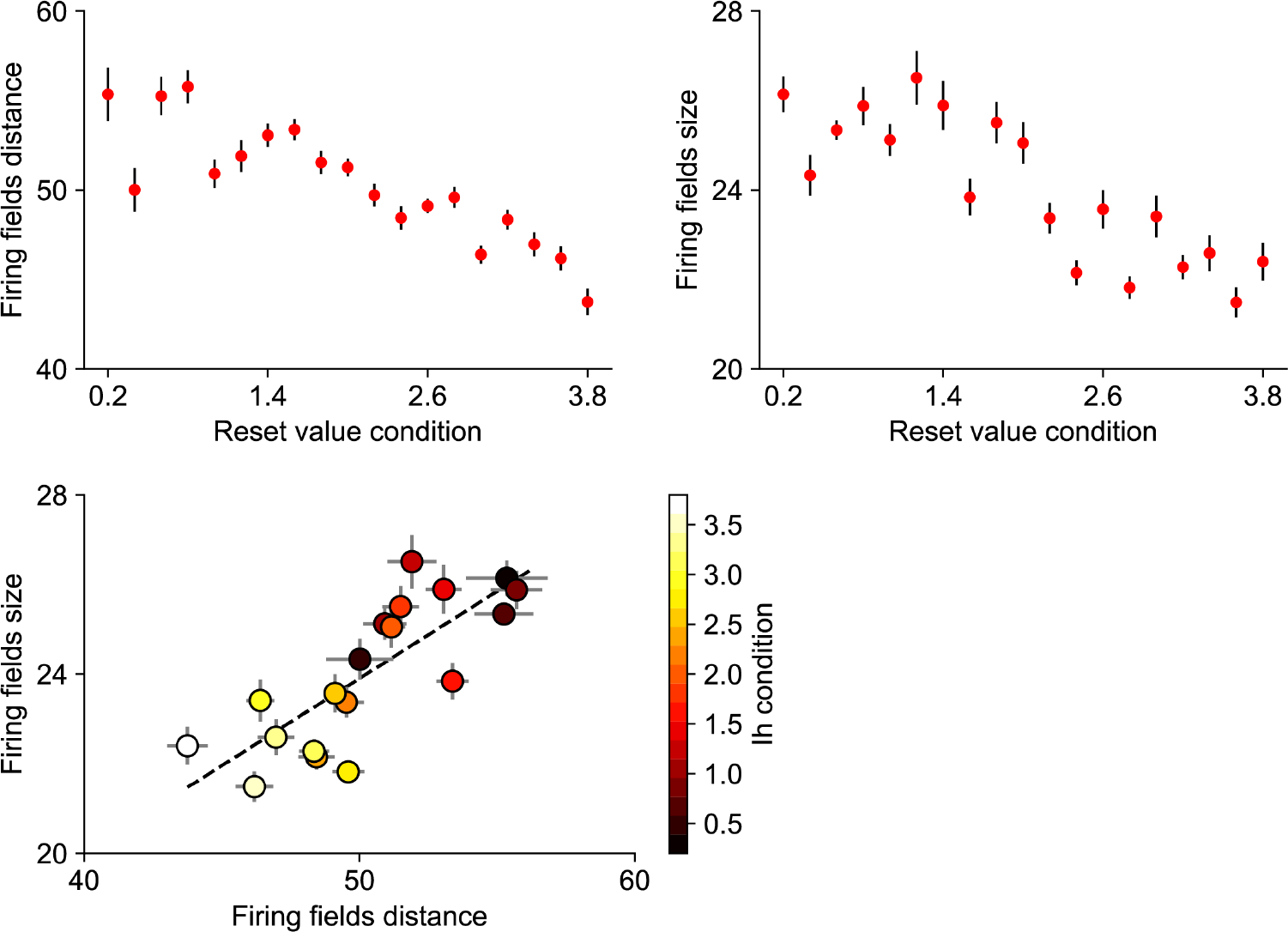
Grid field sizes and spatial distance in simulations run with varying after-spike-reset values. Top: Firing field distance (left) and size (right) decrease along the simulated conditions for larger *d*. Data points represent the average ±SD of all cells in a simulated neuronal population.

Further, we analyzed the relationship between the size and the distance of the firing fields of every cell used in our simulations with the hyperpolarization reset value of each condition through a generalized linear model (GLM). According to the output, firing field size was modeled accordingly by: *logit*(*π*_*i*_) = 5.3 − 0.1 * *size*, and *variance* = 0.019. Firing field distance was modeled accordingly by: *logit*(*π*_*i*_) = 6.6 − 0.07 * *distance*, and *variance* = 0.012.

#### 3.1.2. 2D arena simulation

With the linear track simulations, we have shown that the intrinsic properties of grid cells can effectively modulate firing field size and spacing. However, testing such grid cell properties in a linear trajectory could fail to demonstrate possible deformations in the characteristic grid pattern.

In order to observe the stereotypical pattern of grid cell and the accounts of after-spike hyperpolarization behavior in grid resolution, we have set the virtual agent to perform random exploration within a two-dimensional open field arena. Similar to the 1D runs, we recorded the membrane potentials of stellate cells belonging to different simulated dorsal-to-ventral axis levels. As expected, the firing properties of cells at the ventral level presented larger and further distributed firing-fields when compared with the ones at the dorsal level (Fig. 4)

**Figure 4:**
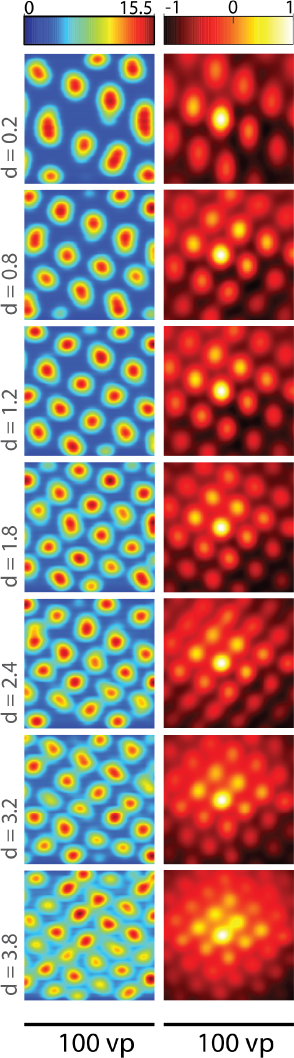
Single cell’s rate maps and autocorrelograms of neurons along dorsal-to-ventral axis. Cell’s spatial activity from ventral (top) to dorsal (bottom) axis level. Hyperpolarization-reset value represented by *d* at left most side. Progressively decrease of grid cell scale (left column), accompanied by its autocorrelogram (right column).

In order to quantify the stability of our model in the grid cells’ spatial representation, we used the gridness score measure for every cell’s autocor-relogram using correlations of rotational symmetry [25], by comparing the spatial autocorrelation maps to the rotated versions of themselves with 30° rotations as:

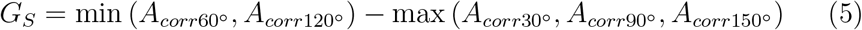

Overall, rate maps along the simulated conditions revealed to be in the range of gridness scores observed in tessellation patterns activity (> 0.15) and were not affected by the hyperpolarization reset value (Pearson *r* = 0.061, *p* = 0.23), suggesting a stable spatial representation across the simulated dorsal-to-ventral axis.

In order to quantify for differences along dorsal-to-ventral levels, we have correlated membrane potentials of cells within each module. Lags calculated after correlating membrane potential signals were taken as a measure of the periodic increase in the amplitude of cells firing rate. Thus, high-resolution grid cell rate-maps at the dorsal level (smaller scale) should reveal shorter distances between firing fields and larger distances for the ones at the ventral level (larger scale). As expected, we observed a progressive decrease in membrane potential autocorrelations lags as a function of the after-spike parameter (Pearson *r* = −0.94, *p* < 0.01, Fig. 5-right). As grid cell hexagonal tessellation patterns and membrane potentials are not dissociable, we have also quantified spatial lags in between firing fields. As for membrane potentials, lags between spatial firing-fields were larger for smaller hyperpolarization reset values (Pearson *r* = −0.91, *p* < 0.01, Fig. 5-left) as well as for rate maps spatial auto-correlation (Pearson *r* = −0.96, *p* < 0.01, Fig. 5-middle). Fig. S2 further illustrates pairwise distances between spatial observations.

**Figure 5:**
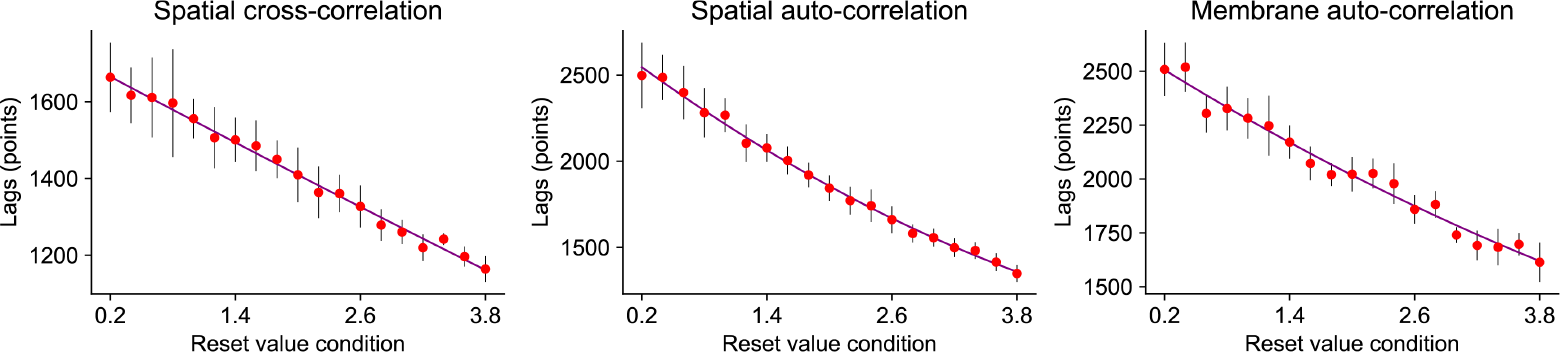
Spatial correlation lags from spike-train and membrane potential along dorsal-to-ventral axis. Lags of correlation along most ventral (0.2) to most dorsal (3.8) conditions. Left and middle plots represent lags of cross- and autocorrelation from 10 cells at each condition. Right plot represents the lags of autocorrelation for membrane potential of each cell. Increase of hyperpolarization-reset value is accompanied by the decrease of lags from ventral to dorsal axis locations.

Despite spatial grid-cell scale distribution found along the dorsal-to-ventral axis levels of MEC layer II, the oscillatory properties of stellate grid cells are also organized on the same axis [10, 18]. To verify whether the hyperpolarization behavior accounting for the spatial resolution organization was also sufficient to modulate the dominant frequencies of simulated neurons, we have compared dominant frequencies of cells at multiple dorsal-to-ventral axis modules. As in [10], dominant frequencies were observed to decrease from dorsal to ventral modules, ranging 14-22 Hz (Pearson *r* = 0.45, *p* < 0.01, Fig. 6-left). Note that frequencies are not in a theta range as expected in MEC, which could be due to the absence of inhibitory projections either from within the MEC population or arriving from hippocampus proper feedback projections. However, there is evidence that modulation of hyperpolarization-after values is sufficient to explain a decrease of membrane potential frequency from dorsal to ventral levels. Thus, spatial scale and oscillatory frequency might be explained by the intrinsic cell hyperpolarization mechanism.

**Figure 6:**
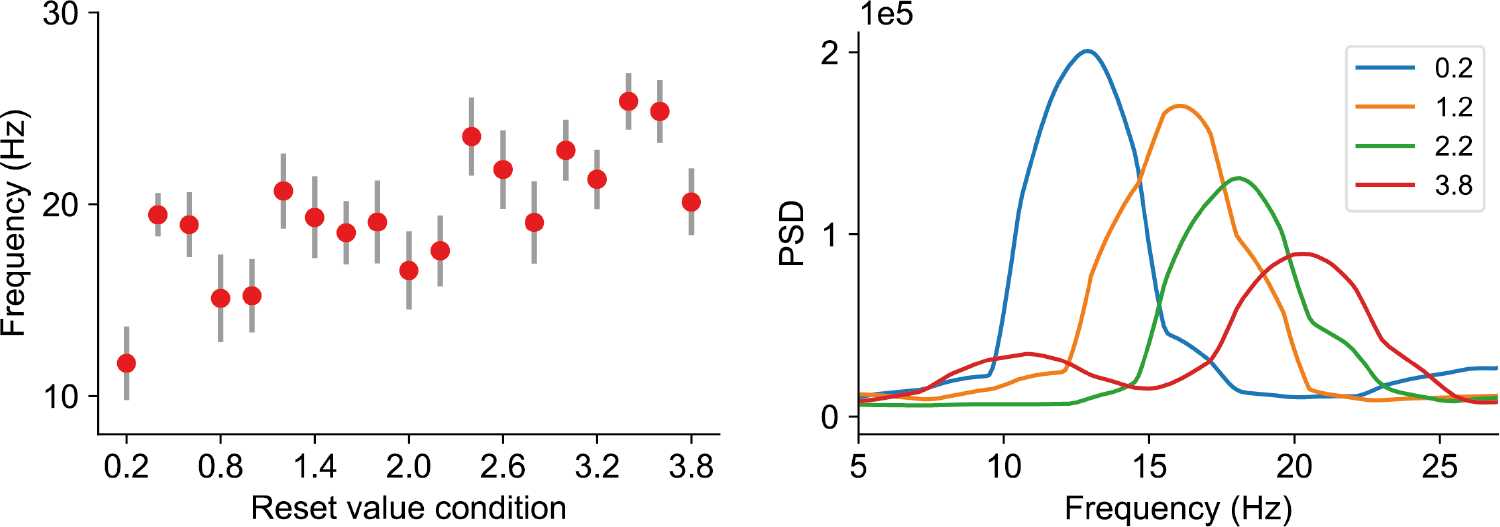
Increase of dominant frequencies from ventral (left) to dorsal (right).

## Discussion

Grid cells in MEC layer II have been characterized by their grid scale, which progressively increases along the dorsal-to-ventral axis. So far, grid cells’ computational models either use oscillatory interference or attractor-based dynamics to elicit the desired behaviors. Hybrid models have also been presented [26]. Regardless, the mathematical formulation to modulate grid scale has been attributed to network dynamics, in the case of attractor-based models, and to network inputs, in the case of interference-based models. Interference models affect its grid cell scale by modulating the frequency of oscillatory signals being projected to the ensemble of grid cells in the network [5]. Attractor dynamics-based grid cell models affect grid cells scale by modulating the gain parameter which reflects how fast the bump of activity in the network is translated to neighbor cells [19]. Despite the fact that both categories of grid cell models use network parameters to affect grid cell scale, it is still unclear what is modulating grid cells’ scale in the hippocampus.

Based on the findings by Giocomo et al. [14, 27], in this study, we hypothesized that grid cells’ scale can be modulated exclusively through single cell properties instead of network properties. To test our hypothesis, we have built upon a previously presented model for grid cells formation based on attractors dynamics [19], as has recently been observed in such cell type [8, 9]. We used spiking neurons to mimic the properties of stellate grid cells and thus modulate their hyperpolarization behavior. We found that after-spike-reset scalars are sufficient to affect both size and scale of grid cells at different axis levels in the medial entorhinal cortex layer II. Specifically, we observed changes in the firing fields’ size and scale, and their respective autocorrelation periodicity bumps from different after-spike-reset values conditions for linear track simulations.

Contrary to Brun et al. [10], we found periodic activity events for the population vector activity analysis. Indeed, one should argue that grid cell periodicity must be noticed at the population level and, thus, whether such a phenomenon is accounted for in a living organism might depend on higher-level spatial encoding mechanisms such as environmental compartmentalization [28].

Because wild rodents typically navigate within two-dimensional environments, we have also tested our library in a virtual agent moving in a 2D arena. As in the 1D environment, the agent had no spatial target position or goal and simply moved at random within the squared arena. Again, the attractor-based network was set to form grid cells as in Guanella et al. [19] using the Izhikevich neuron model [22] to mimic stellate cells found in MEC layer II. The only parameter differing among cells was the after-spike-reset values specific to each sub-population. As expected, size and scale of grid cells firing fields increased progressively along the simulated MEC layer II dorsal-to-ventral axis. In addition, 2D navigation simulations allow us to confirm that gridness remained stable and was not affected by hyperpolarization related properties. Linear decays along simulated conditions were observed for both spatial and membrane potential correlations lag, allowing the quantification of firing-fields distances.

In accordance with Brun et al. [10] and as hypothesized by Navratilova et al. [18], both oscillatory frequencies and spatial scale were affected by cellular after-spike-reset parameters, suggesting that biophysical mechanisms alone are sufficient to modulate multiple grid cell properties.

The flexibility of the synaptic connections has been previously questioned and marked as an implausible mechanism to update the attractor activity bump in the biological brain (McNaughton, 2006). One possibility to over-come such constraint was also discussed in the same paper (McNaughton, 2006), with the solution to rely on multiple networks of conjunctive cells whose activity is dependent on the animal motion. On the other hand, slower mechanisms of synaptic matrix changes might compromise the efficacy of the model‥

We have addressed the question of whether MEC layer II grid cell scale was determined by the network synaptic connectivity distribution, as predicted by low-continuous attractor models, or whether intrinsic properties of stellate cells accounting for individual cell’s hyperpolarization behavior was sufficient to modulate the rate-map resolution of explored environments. Our results suggest that biophysical grid-cell properties are responsible for their spatial scale.

Despite the computational evidence, it is still not clear, however, what are the mechanisms determining the differences in cells response along the dorsal-to-ventral axis level. During development, dorsal regions are formed earlier than the ones at ventral levels [29]. Similarly, as the animal’s development advances, so does its spatial exploration, covering bigger regions of the environment. Thus, it is still uncertain as to what are the causal relation-ships between behavioral components of exploration, such as the magnitude of environmental exploration, and cellular development and organization. In this line, the work of [14] presented differences in the frequency of subthreshold membrane potential oscillations in entorhinal cells. Moreover, in [13], the authors observed a modulation of the cell’s spatial scale in nucleotide-gated (HCN) channels knockout mice compared to sham. Suggesting the functional role of HCN in mediating the topographic organization of firing fields in the explored environment.

Despite the fact that only excitatory cells are used in our implementation, the low-continuous attractor mechanism defines the synaptic weight between cells accordingly to their Cartesian distance in the neural sheet in a range from strong excitation to neighboring cells to strong inhibition to further apart cells. Excitation and inhibition projections from the same neuron is definitely implausible in biological brains, however, that could be solved computationally by setting the synaptic weights to the excitatory range and adding a population of inhibitory interneurons mediating neuronal competition, as suggested by the E%-max winner-take-all mechanism of gamma frequencies (de Almeida et al., 2009)

Along those lines, our work proposes specific physiological and developmental questions that could be tested experimentally. Specifically, characterization of the bursting behavior observed in entorhinal stellate cells along the dorsal-ventral axis, as well as optogenetic stimulation modulating the neuron’s oscillatory dynamics could, potentially, support our modeling results. Moreover, at the computational level, this study proposes future work to unveil the interactions between attractor dynamics and intrinsic properties of stellate cells in the MEC layer II.

## Acknowledgments

*Funding* The research leading to these results has received funding from the European Research Council under the European Union’s Seventh Frame-work Programme (FP7/20072013)/ERC grant agreement no (341196) cDAC.

## Supporting information

**Figure S1:**
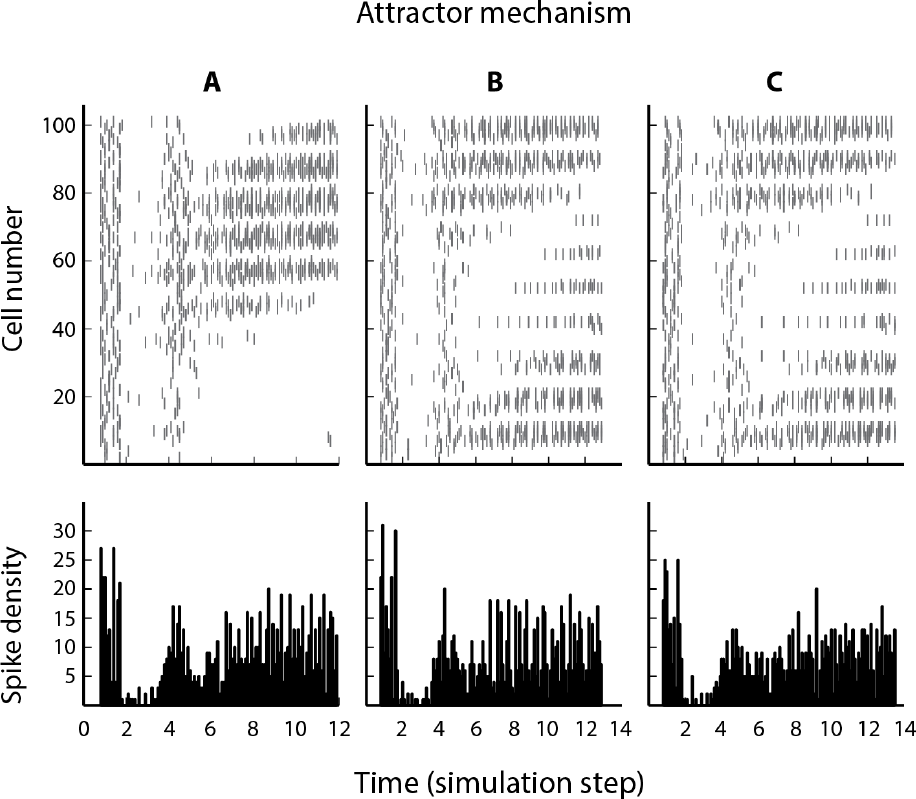
Evidence of attraction at the simulation initial steps. Every cell at each population (assemble) starts with random activity. The bump of activity is formed and attracted to a set of cells. A, B and C represents activity from population 1, 10 and 19, respectively.

**Figure S2:**
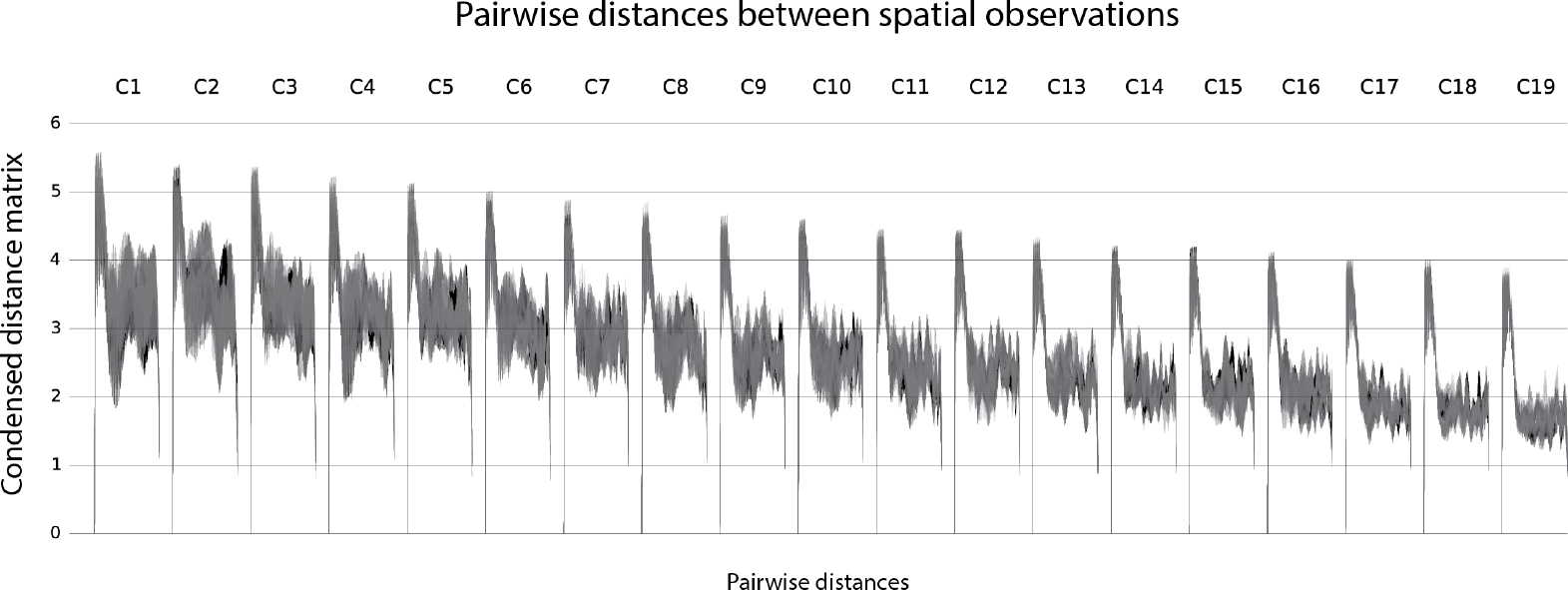
Pairwise distances between spatial observations. Gaussian kernel sigma of firing fields for ten cells at each condition is shown. Decrease of condensed distance matrix against pairwise distances from ventral (left) to dorsal (right) conditions.

